## Supplemental Figures S1-S5 for "Measuring the burden of hundreds of BioBricks defines an evolutionary limit on constructability in synthetic biology"

*These authors contributed equally to this work

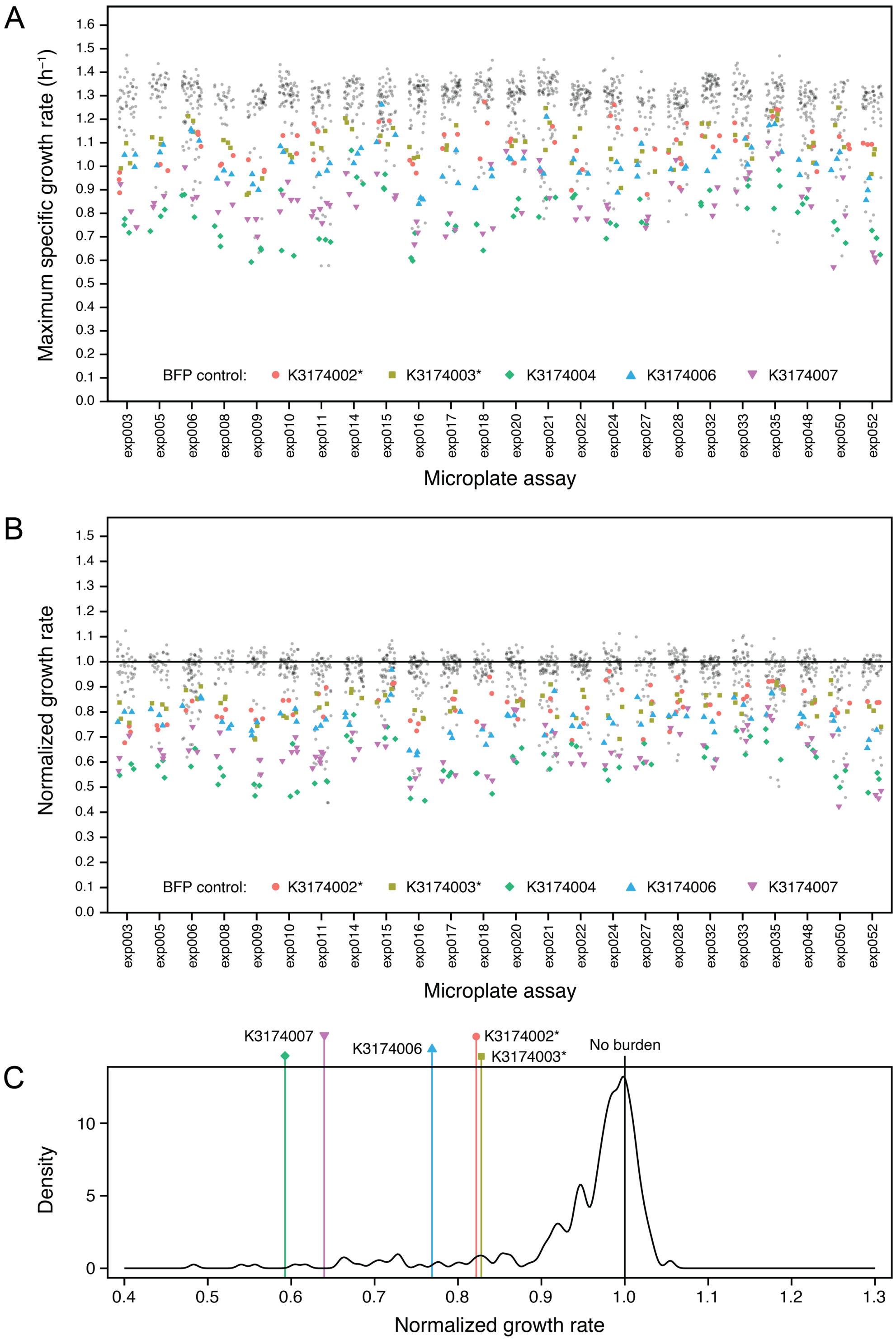


**Fig. S1. Growth rate measurements for all microplate assays.** (**A**) Growth rates fit for each well containing an *E. coli* strain transformed with a BioBrick plasmid across 24 microplate assays. The five highlighted BioBricks are the BFP controls that were included in each assay. Cell stocks of the two starred BFP controls used in these assays had mutations that lowered their burden (**Fig. S3**). (**B**) Normalized growth rates after correcting for variation between assays. (**C**) Final distribution of the mean normalized growth rates estimated for each BioBrick plasmid. The density is graphed using a Gaussian kernel with a bandwidth of 0.005.

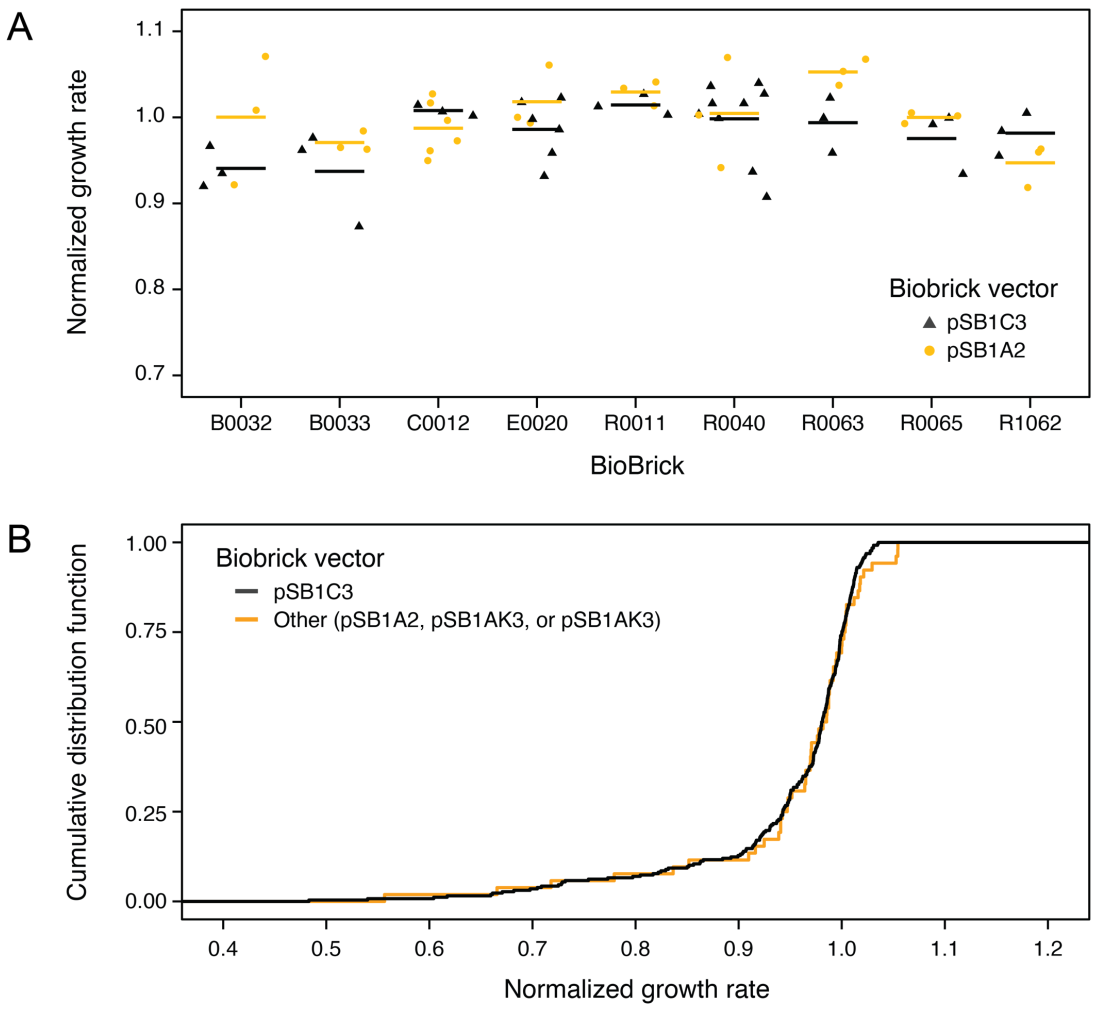


**Fig. S2. Comparison of growth rates measured for BioBricks in different plasmid backbones.** (**A**) Growth rates fit for each well containing an *E. coli* strain transformed with one of the nine BioBricks that were tested in both the pSB1C3 and pSB1A2 backbones. Horizontal bars are mean values. (**B**) Cumulative distributions of the mean normalized growth rate values determined for the 259 BioBricks measured in the pSB1C3 backbone and the 52 measured in other backbones (40 in pSB1A2, 2 in pSB1AK3, and 1 in pSB3C5), excluding the BFP controls.

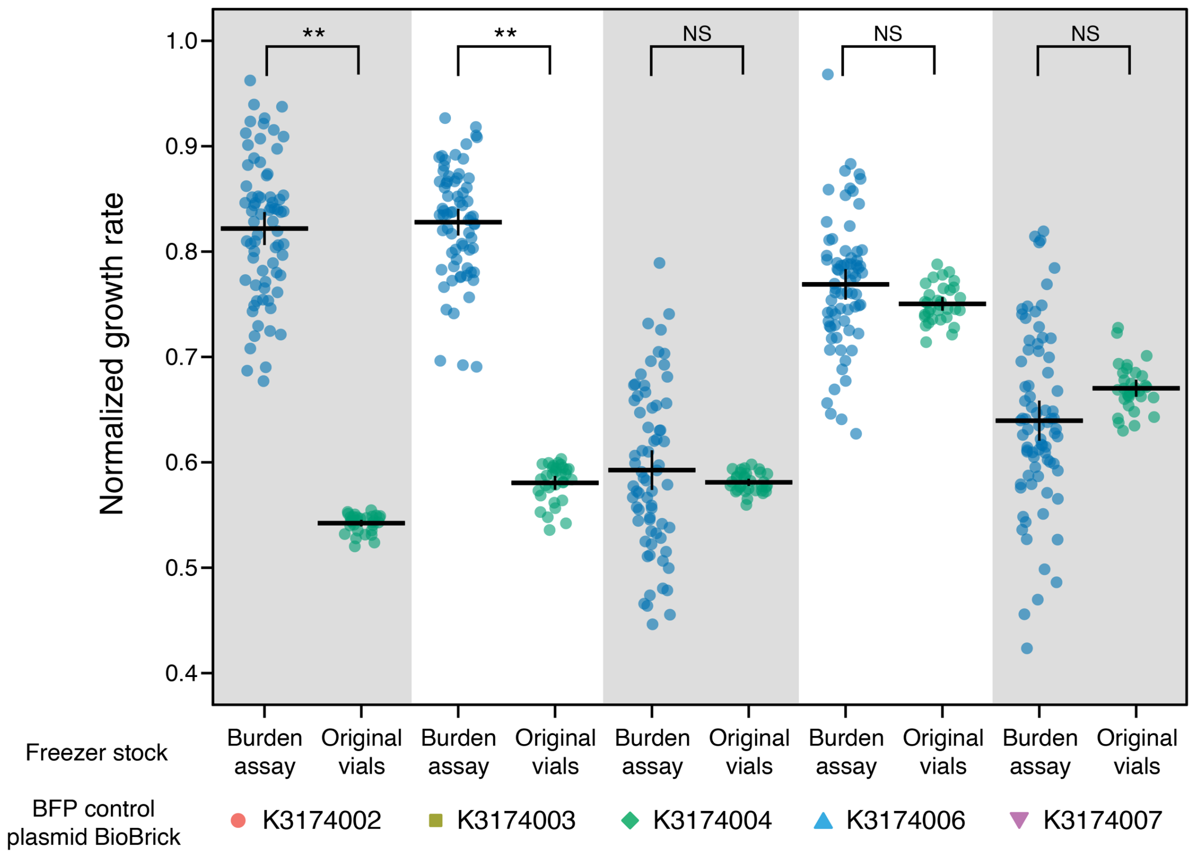


**Fig. S3. BFP plasmids in cell stocks used for microplate assays mutated to reduce burden.** Blue points on the left in each set are normalized growth rate measurements for strains with BFP control strains containing the specified BioBrick plasmids from the freezer stocks used in all of the microplate burden assays (**Fig. S1**, **Fig. S5**). Green points on the right in each set are growth rate measurements of the BFP plasmid control strains grown from the original freezer vials of cells transformed with these plasmids, from which the stocks used in the burden assays were derived through picking colonies and regrowth. Measurements of the original freezer stock strains were normalized to the burden assay normalized growth rates using a linear regression that included only BFP control strains with BioBrick plasmids K3174004, K3174006, or K3174007. The stocks of cells with BioBrick plasmids K3174002 and K3174003 that were used in the burden assays were mutated in a way that greatly reduced their burden. Only the differences between freezer stocks for these two strains had normalized growth rates that were significantly different between the two sets (Bonferroni-corrected two-tailed *t*-tests; ** $p < 0.01$; NS, not significant, $p > 0.05$). Horizontal bars in each set of measurements show means and vertical bars show 95% confidence limits.

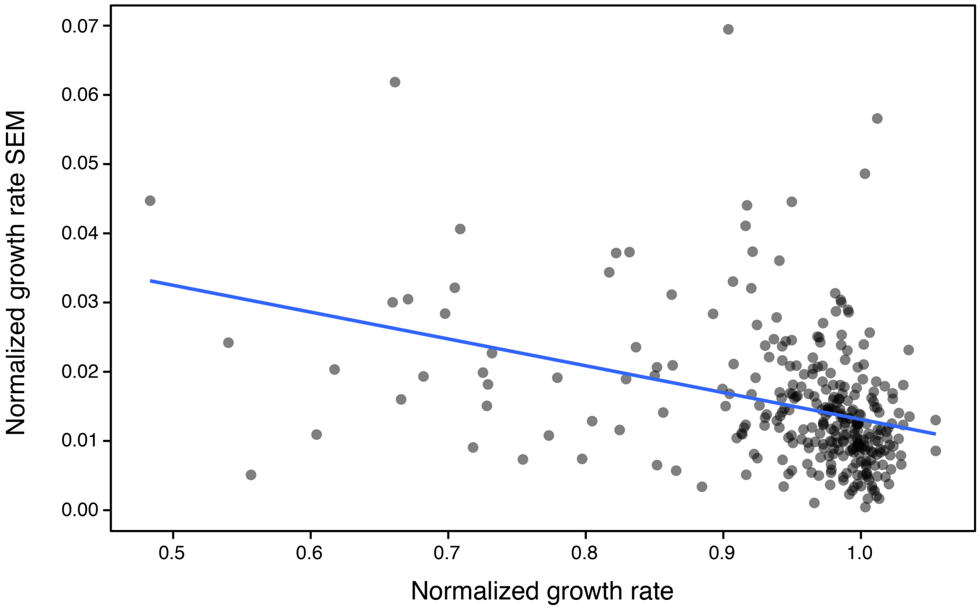


**Fig. S4. Growth rate measurements for BioBricks with higher burden exhibit more variability.** The standard error of the mean (SEM) of the normalized growth rate measured for each of the 301 *E. coli* strains transformed with a different BioBrick plasmid is plotted versus its mean normalized growth rate. The blue trendline is the best-fit least-squares regression. Its slope is significantly different from zero ($p$ = 2.0×10$^{-11}$, two-tailed $t$-test). Strains with lower growth rates (which carry plasmids with higher burden) exhibit more variation in their growth rates across replicate assays.

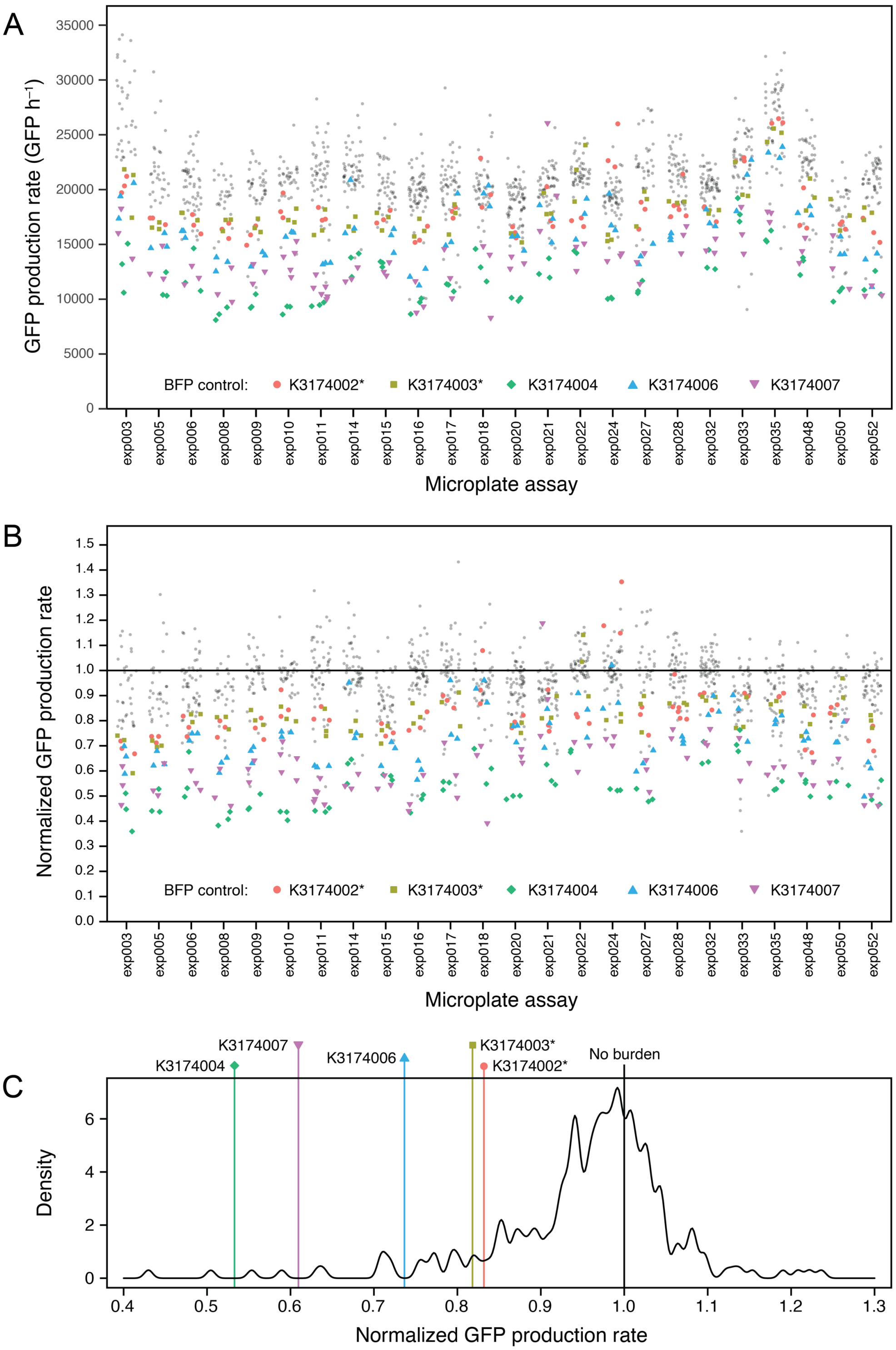


**Fig. S5. GFP production rate measurements for all microplate assays.** (**A**) GFP production rates fit for each well containing an *E. coli* strain transformed with a BioBrick plasmid across 24 microplate assays. The five highlighted BioBricks are the BFP controls that were included in each assay. Cell stocks of the two starred BFP controls used in these assays had mutations that lowered their burden (**Fig. S3**). (**B**) Normalized GFP production rates after correcting for variation between assays. (**C**) Final distribution of the mean normalized GFP production rates estimated for each BioBrick plasmid. The density is graphed using a Gaussian kernel with a bandwidth of 0.005.
